# Identification of disease resistance genes from a Chinese wild 1 grapevine (*Vitis davidii*) by analysing grape transcriptomes and transgenic Arabidopsis plants

**DOI:** 10.1101/169631

**Authors:** Ying Zhang, Jia-Long Yao, Hu Feng, Jianfu Jiang, Xiucai Fan, Haisheng Sun, Yun-Fei Jia, Yanyan Zhu, Ran Wang, Chonghuai Liu

## Abstract

**Highlight:** Transcription profiles showed that 20 candidate genes were obviously co-expressed at 12 hpi in 30 *Vitis davidii*.

VdWRKY53 trancription factor enhanced the resisitance in grapevine and Arabidopsis.

**Abstract:** The molecular mechanisms underlying disease tolerance in grapevines remain uncharacterized, even though there are substantial differences in the resistance of grapevine species to fungal and bacterial diseases. In this study, we identified genes and genetic networks involved in disease resistance in grapevines by comparing the transcriptomes of a strongly resistant clone of Chinese wild grapevine (*Vitis davidii* cv. Ciputao 941, *DAC*) and a susceptible clone of European grapevine (*Vitis vinifera* cv. Manicure Finger, *VIM*) before and after infection with white rot disease (*Coniella diplodiella*). Disease resistance-related genes were triggered in *DAC* approximately 12 hours post infection (hpi) with *C. diplodiella*. Twenty candidate resistant genes were co-expressed in *DAC*. One of these candidate genes, *VdWRKY53* (GenBank accession KY124243), was over-expressed in transgenic *Arabidopsis thaliana* plants and was found to provide these plants with enhanced resistance to *C. diplodiella*, *Pseudomonas syringae* pv *tomato* PDC3000, and *Golovinomyces cichoracearum*. This result indicates that *VdWRKY53* may be involved in nonspecific resistance via interaction with fungal and oomycete elicitor signals and the activation of defence gene expression. These results provide potential gene targets for molecular breeding to develop resistant grape cultivars.

## Introduction

The grapevine is an important fruit crop grown worldwide, and its cultivars are mostly derived from the European species *Vitis vinifera*, which possesses genes for high fruit quality and adaptation to a wide variety of climatic conditions. However, *V. vinifera* cultivars are susceptible to many pathogens, such as phytoplasmas, viruses, bacteria, and fungi (Ferreira *et al.*, 2004).

Infection of grapevines with the fungus *Coniella diplodiella (*Speg.) causes a devastating white rot disease resulting in partial to total crop losses. This disease also has severe impacts on the environment because repeated fungicide applications are required to control the disease. *C. diplodiella* obtains nutrients from infected leaf and berry tissues and eventually decays these tissues. In contrast to *Plasmopara viticola* (causing downy mildew in grapevine) infection with haustoria, *C. diplodiella* exhibits an ambiguous infection pattern. To ward off these tenacious pathogens, plants have developed a vast array of immune responses. Plants, including *Vitis* species, show different resistance levels depending on their different immune mechanisms. In contrast to *V. vinifera* cultivars, most clones of the Chinese wild species *DAC* exhibit high levels of resistance (Ferreira *et al.*, 2004), allowing major quantitative trait loci (QTL) of resistance genes (Bellin *et al.*, 2009; Marguerit *et al.*, 2009) to be mapped. QTL mapping will assist breeders in the introgression of these genes into *V. vinifera*-based cultivars. Conventional breeding has created some interspecific hybrids that are resistant to fungal diseases, but this process is long and inefficient. Further breeding work is required to combine resistance with berry quality suitable for table and wine grapes. This breeding process will be greatly accelerated by the availability of grapevine genome sequences (Jaillon *et al.*, 2007) and marker-assisted selection (Jaillon *et al.*, 2007; Costantini *et al.*, 2008).

Of approximately 70 *Vitis* species worldwide, 38 originate in China (Kong, 2004). Chinese wild grapevines are a very important source of grapevine germplasm for breeding new cultivars. Studies have already revealed that Chinese wild grapevines possess resistance genes and special resistance mechanisms (Li *et al.*, 2012). We have collected 500 accessions from 20 Chinese wild grapevine species in a germplasm nursery and found one accession, *Vitis davidii* 0941, with the highest level of resistance after *in vitro* testing (Zhang *et al.*, 2013). However, it is not yet known which genes regulate these resistance traits in grapes.

The expression of resistance genes is often induced by pathogen infection in resistant plants (Tao *et al.*, 2003; Tripathi *et al.*, 2012; Phukan *et al.*, 2016). This type of expression may be much weaker or even absent in susceptible plants. Resistance genes are among the differentially expressed genes (DEGs) that can be identified by comparative analysis of the transcriptomes of resistant and susceptible plants after infection (Ma *et al.*, 2016). Therefore, we chose to use this technology to reveal the DEGs between susceptible and resistant grapevine species associated with resistance responses to *C. diplodiella* from early to late stages of infection.

We found that *Vitis davidii* expressed resistance genes in the early stage of infection, but *V. vinifera* did not. We further identified 20 candidate resistance genes. Over-expression of one of the candidate genes, *VdWRKY53*, in transgenic Arabidopsis plants conferred resistance to *C. diplodiella*, *Pseudomonas syringae* pv *tomato* PDC3000, and *Golovinomyces cichoracearum* (powdery mildew of Arabidopsis).

## Materials and methods

### Observations of microstructure and ultrastructure of leaves

Leaf samples 0.25 cm^2^ in area were collected from the middle of five healthy mature leaves from *Vitis vinifera* cv. Manicure Finger (*VIM*) and *Vitis davidii* cv. Ciputao 0941(*DAC*)and fixed in FAA fixative (mixing alcohol acetate formalin fixative). After the leaves became transparent, they were washed in water for 4 h, transferred into a solution of glycerol: lactic acid: water (volume ratio 1: 1: 1) for 24 h and stored until use. The leaves were stained for 20 min in 0.5% aniline blue (dissolved in a mixture of glycerol: lactic acid: water, volume ratio 1: 1: 1), rinsed with ethanol (Vanacker *et al.*, 2000) and observed under a dissecting microscope.

The samples were then embedded in paraffin and cut into 10-μm-thick sections that were stained with haematoxylin and then examined under a microscope (Lycra DMi1-PH1, Germany) and photographed with an Olympic BX51 camera, Japan (Lighezan *et al.*, 2009).

The ultrastructure was observed using a transmission electron microscope (TEM) based on the method described by Lighezan *et al*. (2009). Images were taken with a HITACHI 7000 TEM.

### Plant material and pathogen inoculation treatments on grapevine

For mRNA sequencing analysis, *Vitis vinifera* cv. Manicure Finger (VIM) and *Vitis davidii* cv. Ciputao 941 (DAC) were grown in a greenhouse at 28°C with a 16 h photoperiod and inoculated with *C. diplodiella* (Speg.) (strain WR01, from the Institute of Plant Protection, CAAS) mycelium gelose discs from a 7-day-old culture grown at 28°C on potato dextrose agar (PDA) medium. Leaf samples were collected at 0, 12 and 36 hpi and were immediately frozen in liquid nitrogen and stored at -80°C in a freezer. Each sample consisted of pooled specimens from three leaves of three plants.

### RNA extraction, library construction, and RNA-seq

For RNA-seq library construction, the total RNA was extracted from six samples using the Total RNA Extraction Kit (BioFlux, Tokyo, Japan). The samples were named *VIM1*, *VIM2*, *VIM3*, *DAC1*, *DAC2*, and *DAC3* for *VIM* and *DAC* collected at 0, 12 and 36 hpi, respectively. The RNA quality and purity were checked using 1% agarose gels and a NanoPhotometer^®^ spectrophotometer (IMPLEN, CA, USA).

Sequencing libraries were generated using 3 μg of RNA per sample as input material and the NEB Next^®^ Ultra™ RNA Library Prep Kit for Illumina^®^ (NEB, USA). Index codes were added to each sample to tag its sequences. Library quality was assessed on an Agilent Bioanalyzer 2100 system. The index-coded samples were clustered on a cBot Cluster Generation System using the TruSeq PE Cluster Kit v3-cBot-HS (Illumina) according to the manufacturer’s instructions. After cluster generation, the libraries were sequenced by Novogene (Beijing) on an Illumina HiSeq 2000 platform to generate 100-bp paired-end reads.

### Data assembly and analysis

Raw data (raw reads) in fastq format were first processed using in-house Perl scripts. Clean data (clean reads) were obtained by removing reads containing the adapter, reads containing poly-N and low-quality reads from the raw data. At the same time, the Q20, Q30 and GC contents of the clean data were calculated. All downstream analyses were based on the clean data with high quality.

Reference genome and gene model annotation files were downloaded from the genome website (http://plants.ensembl.org/Vitisvinifera). An index of the reference genome was built using Bowtie v2.0.6, and paired-end clean reads were aligned to the reference genome using TopHat v2.0.9. The number of reads mapped to each gene was counted using HTSeq v0.5.4p3. In addition, the RPKM (Reads Per Kilobase of transcript per Million mapped reads) of each gene was calculated based on the length of the gene transcript, the reads mapped to the gene, and the total mapped reads (Mortazavi *et al.*, 2008).

Prior to differential gene expression analysis, for each sequenced library, the read counts were adjusted by the edge R program package with one normalized scaling factor. The differential expression under two conditions was analysed using the DEGseq R package (1.12.0). The P values were adjusted using the Benjamini & Hochberg method. A corrected P value of 0.001 and a log2 (Fold change) of 1 were set as the threshold for significantly differential expression.

### Gene co-expression analysis

Gene co-expression network analysis was performed on each RNA-seq library to analyse the correlation of genes from each experimental sample, followed by a search for resistance-related pathways and genes (Gillis and Pavlidis, 2011). We also used the WGCNA method to test the effect of thresholding the networks (Zhang and Horvath, 2005). We subjected the best WGCNA results to MeV K-means analysis, setting the cluster number to 50 (k=50).

### Annotation and functional classification

Gene Ontology (GO) enrichment analysis of the DEGs was implemented with the GO seq R package, in which gene length bias was corrected. GO terms with corrected P values less than 0.05 were considered to be significantly enriched in the DEGs. We used the KOBAS software to test the statistical enrichment of the differential expression of genes in KEGG pathways. Putative gene functions were assigned using a set of sequential BLAST searches of all the assembled unigenes against sequences in the Ensembl Plants (http://plants.ensembl.org/Vitisvinifera) database of non-redundant proteins and nucleotides, the Swiss-Prot protein database, the Gene Ontology database, the Cluster of Orthologous Groups database, and the Kyoto Encyclopedia of Genes and Genomes database.

### qRT-PCR analysis

The total RNA of grapevines was extracted using the improved SDS/phenol method as described by (Ülker *et al*. 2007). PCR primers for the reference gene *EF1 ｒ* and the test genes are listed in Table S1. Three independent PCR reactions were conducted for each gene using a LightCycler^®^ 480 (Roche Diagnostics, Rosel, Switzerland), and the relative expression levels of the genes were calculated using the 2^-_ΔΔ^ct^_^ method (Ramamoorthy *et al.*, 2008).

### Vector construction and Arabidopsis transformation

The full-length cDNA of *VdWRKY53* was amplified by PCR and cloned into the BglII/BstE2 site of the binary plasmid pCAMBIA3301, generating pCAMBIA3301-VdWRKY53 (pGW53). This new plasmid was verified by sequencing and then introduced into *A. tumefaciens* GV3101 cells for Arabidopsis transformation via the floral dipping method (Clough and Bent, 1998).

### Pathogenic fungus/bacterium inoculation on Arabidopsis

*C. diplodiella* was tested on wild-type (Columbia (Col)) and transgenic (GW53) Arabidopsis plants, which were grown in a culture chamber (25°C; 12 h photoperiod; light intensity 100 μmol m^-2^s^-1^). The powdery mildew (*Golovinomyces cichoracearum*) isolate UCSC1 was cultured on Arabidopsis *phytoalexin cichoracent 4* (*pad4*) mutant plants. Powdery mildew inoculation of Arabidopsis was performed as previously described (Wang *et al.*, 2007).The bacterial strain *Pseudomonas syringae* pv tomato PDC3000 was grown in LB liquid medium (Yu *et al.*, 2011), and the inoculation method was based on Melotto *et al*. (2008).

## Results

### *Anatomical structure of* Vitis *leaves and inoculation in grapevines*

*DAC* is an important wild grapevine species that grows in 10 provinces in China. In our experiments, *DAC* showed the highest level of resistance among all grapevines tested (Zhang *et al.*, 2013; *Zhang and Feng, 2014*). Examination of the anatomical structure revealed that the leaves of *DAC* were not significantly different from those of *VIM* in thickness, including the thickness of the palisade, spongy tissues, and upper and lower epidermis (Table 1, Fig. 1A). Except for a higher number of chloroplasts in the palisade of *DAC* than in that of *VIM* (Fig. 1A), which is important in photosynthesis, *DAC* had a similar leaf structure to *VIM*. This result suggests that the differences in disease resistance between *DAC* and *VIM* are unlikely to be due to the differences in their leaf structure and more likely to be due to the differences in their resistance genes.

**Table 1.** Comparison of leaf tissue thickness of two Vitis species.

| Species | Leaf thickness<br>($\mu\text{m}$) | Upper epidermis<br>thickness ($\mu\text{m}$) | Palisade tissue<br>thickness ($\mu\text{m}$) | Spongy tissue<br>thickness ($\mu\text{m}$) | Lower epidermis<br>thickness ($\mu\text{m}$) |
| --- | --- | --- | --- | --- | --- |
| *Vitis davidii* | $91.13 \pm 7.48^a$ | $12.43 \pm 2.15^a$ | $36.44 \pm 4.32^a$ | $36.33 \pm 7.70^a$ | $7.87 \pm 1.72^a$ |
| *Vitis vinifera* | $112.78 \pm 18.50^a$ | $12.56 \pm 2.82^a$ | $42.75 \pm 8.53^a$ | $48.98 \pm 8.49^a$ | $9.43 \pm 1.84^a$ |

\* Data are mean ± sd, n = 12, significant difference by analysis of variance.

**Fig. 1.**
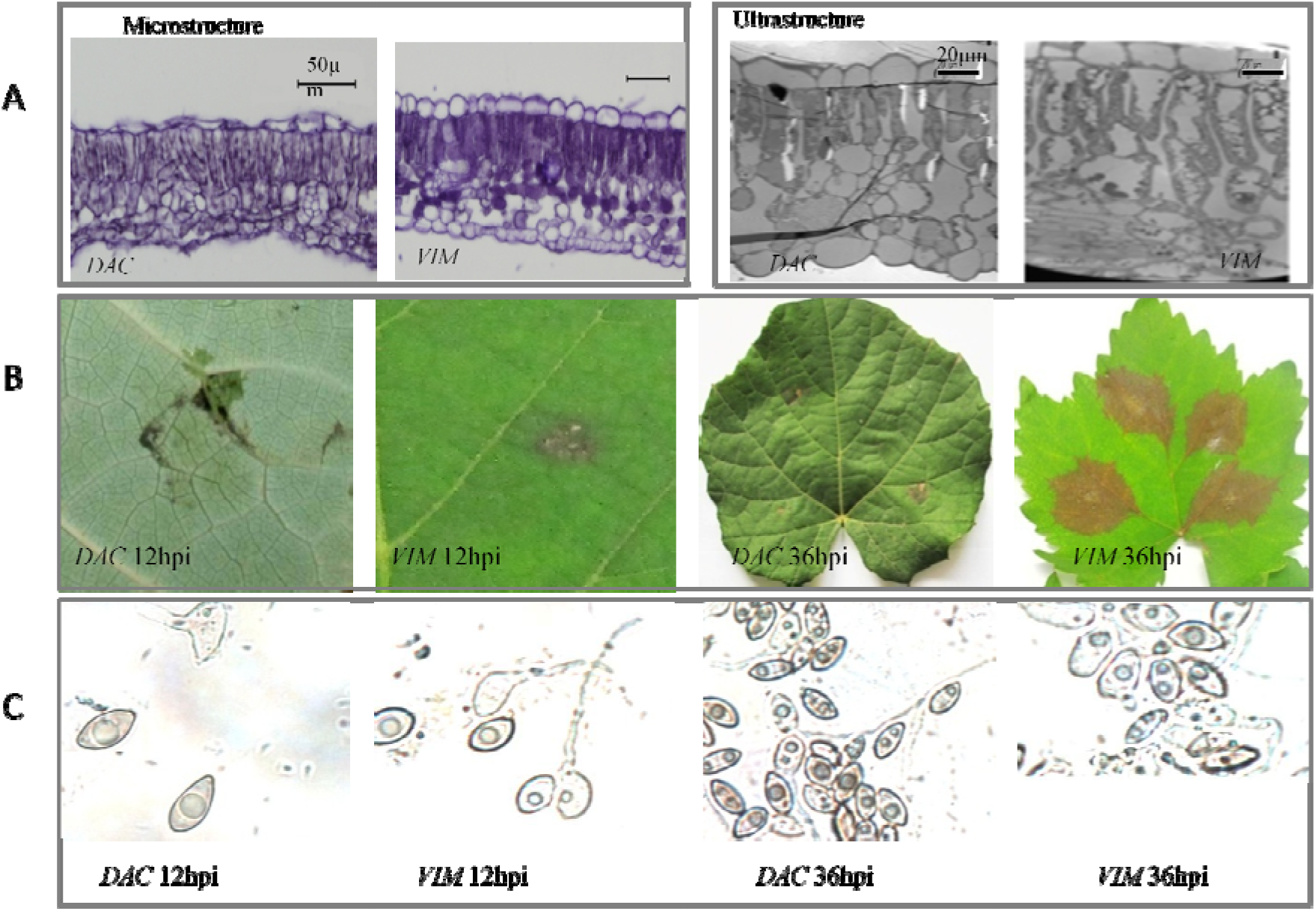
Vitis leaf structure and symptoms of *C. diplodiella* infection. (a) Anatomical structure of in *Vitis vinifera* cv. Manicure Finger (*VIM*) and *Vitis davidii* cv. Ciputao 0941(*DAC*). Leave samples were collected from 2-week old leaves at the 3-4 position on a branch. At microstructure level, leaf thickness, upper epidermis thickness, palisade tissue thickness, spongy thickness, lower epidermis thickness were not significant variation in between *DAC* and *VIM*. At ultrastructure level, *DAC* had more chloroplasts in palisade than *VIM.* (b) Symptoms in *DAC* and *VIM* after *C. diplodiella* infection. Typical hypersensitive response (HR) symptoms were shown in *DAC* but not in *VIM* at 12 hpi (hours post infection) and 36 hpi. (c) *C. diplodiella* infected leaves were examined under a microscope at 12 hpi and 36 hpi to show fungal spore germination and growth. *C. diplodiella* spore germination was observed on *VIM* leaves at 12 hpi and on *DAC* leaves at 36 hpi.

After infection with *C. diplodiella*, the *DAC* leaves showed weaker disease symptoms than those on *VIM* leaves at 12 hours post infection (hpi). At 36 hpi, the symptoms on *DAC* leaves developed into a typical hypersensitive response (HR), where cell death at the infection site blocked further spreading of the pathogen, but the symptoms on *VIM* leaves developed into typical grape white rot disease (Fig. 1B). Under a microscope, germination of spores of *C. diplodiella* was observed on *VIM* leaves but not on *DAC* leaves at 12 hpi. At 36 hpi, spore germination was completed on the leaves of both grape species, but the germination rate was lower on *DAC* leaves than on *VIM* leaves (Fig. 1C). The hyphal growth rate and state were similar on leaves of the two grape species. Given the importance of 12 and 36 hpi in disease development, we chose these two time points for transcriptome comparison with the 0 hpi stage.

### mRNA sequencing statistics

To analyse the transcript levels of *DAC* and *VIM* at 0, 12, and 36 hpi with *C. diplodiella*, approximately 433 million reads were generated from the six libraries (*VIM1*, *VIM2*, *VIM3*, *DAC1*, *DAC2*, and *DAC3*). These reads constitute 42.32 Gb of cDNA sequence (Table 2). Among them, 97.2% were high-quality (Q>20) reads and were selected for further analyses. Of the clean reads, 76.96-88.01% were mapped to the *Vitis vinifera* reference genome (http://plants.ensembl.org/Vitisvinifera) (Jaillon *et al.*, 2007). For each library, the reads were mapped to approximately 23-25 thousand genes, of which approximately one thousand were novel genes that were not annotated in the grape reference genome (Table 2).

**Table 2.** Summary of transcriptomes data.

|  | VIM1 | VIM2 | VIM3 | DAC1 | DAC2 | DAC3 |
| --- | --- | --- | --- | --- | --- | --- |
| Total reads (x1000) | 76,192 | 76,999 | 67,696 | 67,447 | 71,886 | 73,179 |
| Base number (G) | 7.62 | 6.7 | 6.76 | 6.74 | 7.18 | 7.32 |
| High-quality reads (%) | 98.1 | 96.48 | 97.97 | 96.31 | 98.02 | 96.46 |
| Mapped reads (%) | 88.01 | 87.27 | 87.43 | 77.33 | 76.96 | 77.14 |
| Number of transcripts | 23,621 | 24,301 | 24,407 | 2,3441 | 25,537 | 24,018 |
| Number of novel transcripts | 961 | 971 | 997 | 938 | 1,020 | 957 |

### Identification of differentially expressed genes (DEGs)

A total of 12976 genes were selected for DEG analyses on the basis that they had a RPKM value greater than 1 (value >1) according to DEGseq. The number of DEGs between the resistant (*DAC*) and susceptible (*VIM*) grape species at each sampling point was estimated using the criteria P<0.001 and log2 (fold change) >2.0 or <2.0.

After *C. diplodiella* inoculation, we detected a total of 7073 transcripts, exhibiting an increase compared with the 0 hpi time point. More transcripts were induced in DAC than in VIM. 256 transcripts of the specific expression in DAC was of unknown genes showing a large genetic distance from the reference genome (http://plants.ensembl.org/Vitis_vinifera) (Fig. 2).

**Fig. 2.**
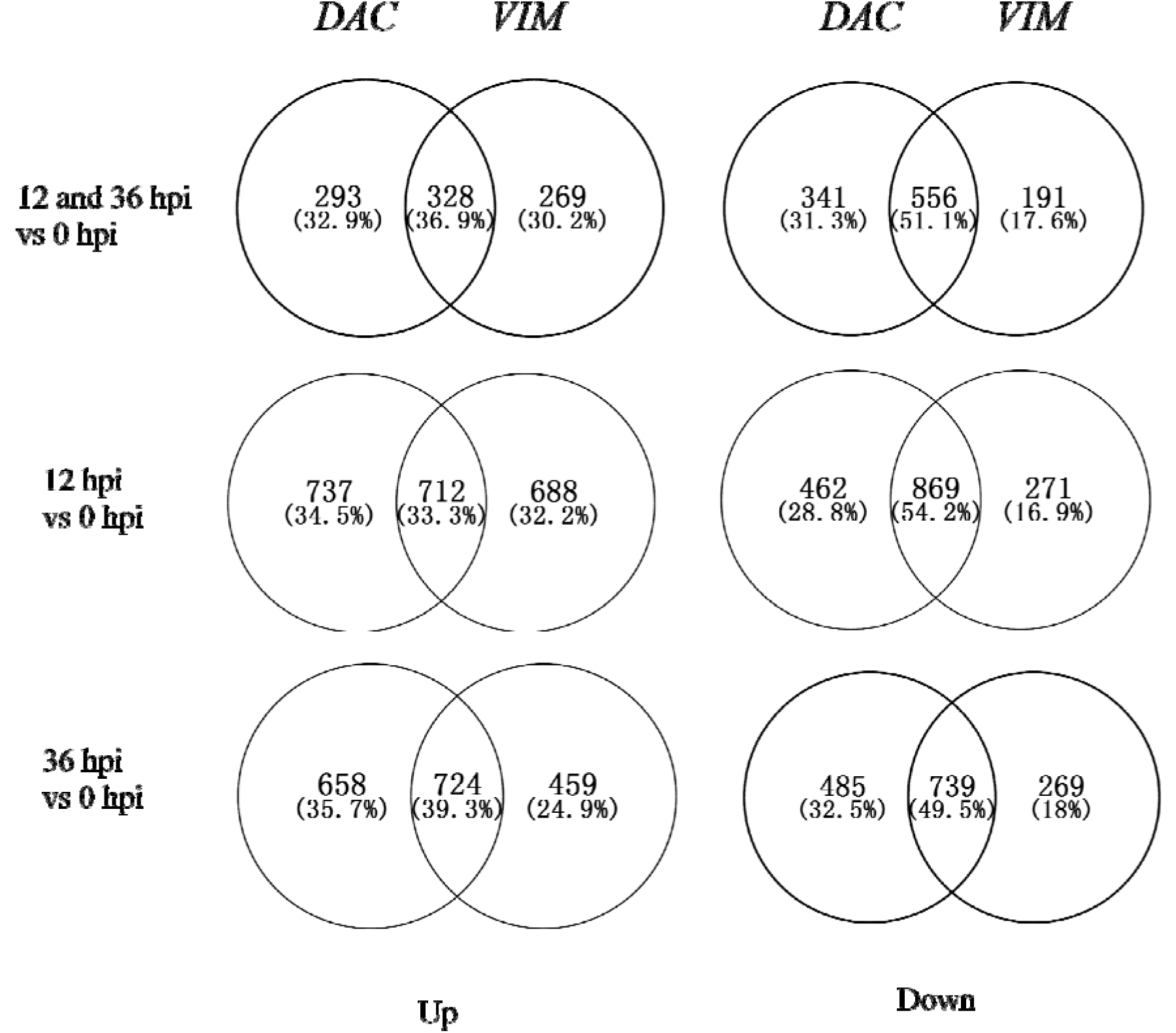
Number of differentially expressed genes (DEGs) in DAC and VIM leaves following infection with *C. diplodiella*. Venn diagrams show the number of DEGs, up or down regulated (P < 0.0001, fold > 2.0), in *DAC* and *VIM* leaves at 12, 36 hpi, and both 12 and 36 hpi compared to 0 hpi. There were more DEGs in resistant genotype *DAC* than in the susceptible genotype *VIM*.

All DEGs were annotated. COG, GO, KEGG, Swiss-Prot and nr were analysed based on their references (Altschul *et al.*, 1997; Tatusov *et al.*, 2000; Kanehisa *et al.*, 2004) (Table 3). To determine the DEGs and pathways between the susceptible *VIM* and the resistant *DAC*, we analysed the KEGG pathways in detail. We found two pathways with no DEGs (Brassinosteroid biosynthesis and Cutin, suberine and wax biosynthesis) and 118 pathways with DEGs. Differences in gene expression at two time points after pathogen infection in *VIM* and *DAC* were examined, and DEGs were identified by pairwise comparisons of the six libraries (Table S2, Table S3).

**Table 3.** Annotation of differentially expressed genes (DEGs) from six libraries.

| DEG Set | Annotated | COG | GO | KEGG | Swiss-Prot | nr |
| --- | --- | --- | --- | --- | --- | --- |
| DAC2_vs_DAC1 | 3,952 | 1,617 | 3,517 | 675 | 2,997 | 3,952 |
| DAC3_vs_DAC1 | 4,025 | 1,657 | 3,594 | 677 | 3,114 | 4,025 |
| VIM2_vs_VIM1 | 3,925 | 1,664 | 3,506 | 656 | 3,031 | 3,925 |
| VIM3_vs_VIM1 | 3,869 | 1,572 | 3,455 | 636 | 2,986 | 3,869 |

### *Candidate genes in resistant grapevine* DAC

We focused on the Plant-pathogen interaction pathway, which involved DEGs including cell wall genes, LRR receptor-like genes, *WRKY* genes and pathogenesis-related (PR) genes. All transcripts were divided into four co-expression groups. Among them, the molecular function group genes were divided into 50 co-expression clusters (K=50) by MeV K-means analysis, corresponding to the RPKM value. The transcript levels of these genes were higher in DAC than in VIM at both stage 0 hpi and 12 hpi, the transcript levels of these genes were reduced at 36 hpi compared to 0 hpi and 16 hpi in both VIM and DAC (Fig. 3 Table S4).

**Fig. 3.**
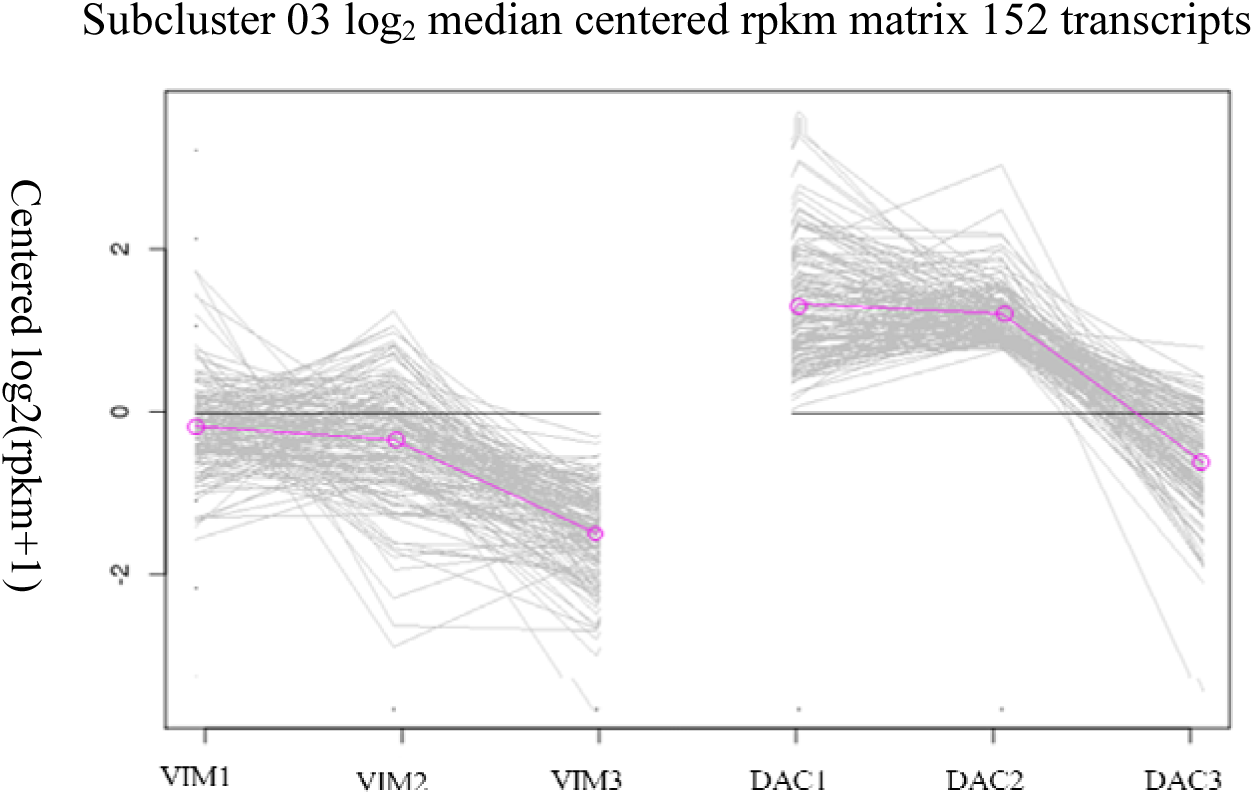
The subcluster 03 from analysis coexpression by K-means analysis. All DEGs were divided 50 co-expression clusters (K=50) by MeV K-means analysis. In Subcluster 03(k=3),152 DEGs were clustered significant coexpression tendency at 12 hpi, red arrow indicate to the expression genes.

**Fig. 4.**
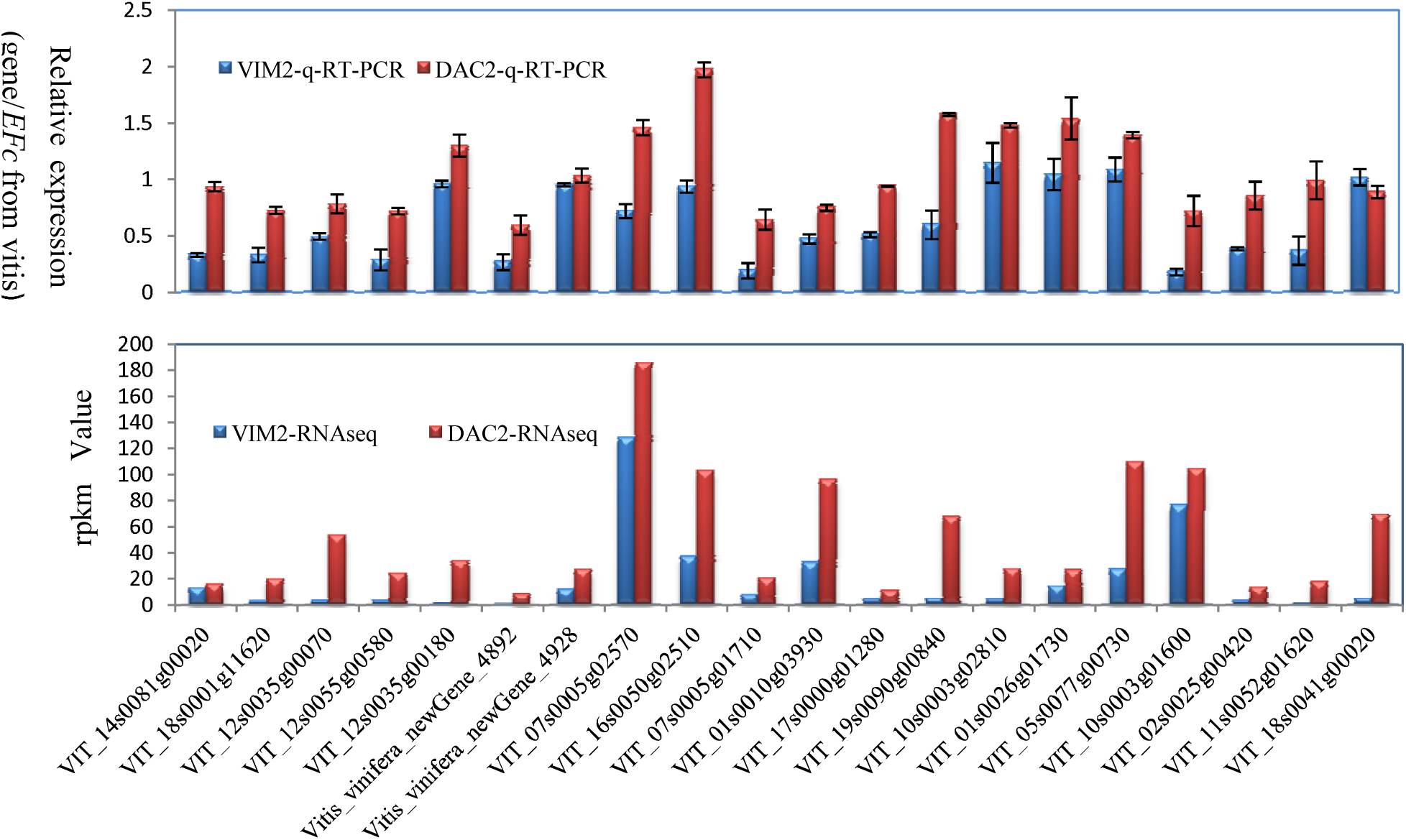
Comparison of RNA-seq data with quantitative RT-PCR results in candidate genes. 152 DEGs were clustered in K03 showed 20 resistance related genes significant different expression at 12 hpi (Figure3). Candidate genes (at 12hpi) identified by both RNA-seq and quantitative RT-PCR, respectively. The numbers 20 genes identified by RNA-seq, which represent Wall-associated receptor kinase 2(VIT_18s0041g00020, VIT_18s0001g11620), LRR receptor-like serine/threonine-protein kinase (VIT_12s0035g00070, VIT_12s0055g00580, VIT_12s0035g00180, Vitis_vinifera_newGene_4892, Vitis_vinifera_newGene_4928), WRKY transcription factors (VIT_07s0005g02570, VIT_16s0050g02510, VIT_07s0005g01710, VIT_01s0010g03930, VIT_17s0000g01280, VIT_19s0090g00840, VIT_10s0003g02810, VIT_01s0026g01730, VIT_05s0077g00730, VIT_10s0003g01600, VIT_02s0025g00420), Pathogenesis-related protein PR (VIT_11s0052g01620, VIT_14s0081g00020)._

Based on the analysis of the transcript expression, which was highly correlated with the resistance response, we suggested 20 candidate genes that work together in the SA signal pathway for *DAC* resistance to *C. diplodiella* and are co-expressed in the K03 cluster. The expression of these genes in *DAC* showed significant differences from their expression in *VIM*. The RPKM measure of read density reflects the molar concentration of a transcript (Mortazavi *et al.*, 2008), We assessed the candidate genes based on two factors, RPKM value (RPKM>1 after inoculation) and the expression change fold (fold>2). There were two wall-associated receptor kinase genes, five LRR receptor-like serine/threonine-protein kinase genes, eleven WRKY transcription factor genes, and two PR protein-like genes in the same resistance signalling pathway. We compared the data between the two species from the six libraries (Table 4). We identified the expression of twenty candidate genes by comparing the RPKM values from the RNA-seq data with the quantitative RT-PCR results at 12 hpi. Nineteen genes were found to have high RPKM values and up-regulated expression, while one gene (VIT_14s0081g00020) with a high RPKM value was down-regulated at 12 hpi in *DAC* (Fig. 3).

**Table 4.** The Co-expression in K03 cluster details of candidate genes involving immune system of DAC were related to resistant to C. diplodiella.

| Gene ID | Gene function | RPKM value |  |  |  |  |  |
| --- | --- | --- | --- | --- | --- | --- | --- |
|  |  | VIM1 | VIM2 | VIM3 | DAC1 | DAC2 | DAC3 |
| VIT_18s0041g00020 | Wall-associated receptor kinase 2 | 9.689896 | 13.35789 | 10.79321 | 22.35743 | 16.1210622 | 76.36481 |
| VIT_18s0001g11620 | Wall-associated receptor kinase 2 | 0.42436 | 2.479711 | 0.601132 | 5.246519 | 19.84848 | 1.520792 |
| VIT_12s0035g00070 | LRR receptor-like serine/<br>threonine-protein kinase | 1.54826 | 2.86169 | 1.56953 | 16.8916 | 53.6436 | 54.9697 |
| VIT_12s0055g00580 | LRR receptor-like serine/<br>threonine-protein kinase | 1.351952 | 2.5520103 | 1.841092 | 5.4257649 | 24.084457 | 5.5218482 |
| VIT_12s0035g00180 | LRR receptor-like serine/<br>threonine-protein kinase | 0.750406 | 0.814589 | 1.07202 | 1.25927 | 33.9365 | 2.5213 |
| Vitis_vinifera<br>_newGene_4892 | LRR receptor-like serine/<br>threonine-protein kinase | 0.899727 | 0.0245324 | 0.04971 | 1.79436 | 8.41789 | 1.21961 |
| Vitis_vinifera<br>_newGene_4928 | LRR receptor-like serine/<br>threonine-protein kinase | 1.67189 | 11.7781 | 0 | 10.7038 | 27.3047 | 5.41412 |
| VIT_07s0005g02570 | WRKY transcription factor | 20.39015 | 128.83849 | 26.54005 | 34.65061 | 186.7816 | 36.9177 |
| **VIT_16s0050g02510** | **WRKY transcription factor** | 15.6605 | 36.7449 | 23.8597 | 21.4326 | 103.314 | 24.7179 |
| VIT_07s0005g01710 | WRKY transcription factor | 6.5624 | 7.35796 | 3.60829 | 6.81616 | 20.64 | 3.8377 |
| VIT_01s0010g03930 | WRKY transcription factor | 4.07527 | 32.4315 | 5.70703 | 22.1046 | 96.5928 | 21.8077 |
| VIT_17s0000g01280 | WRKY transcription factor | 0.306743 | 3.68295 | 0.441533 | 2.62295 | 10.4016 | 9.98668 |
| VIT_19s0090g00840 | WRKY transcription factor | 0.4447659 | 4.63589 | 0.30357971 | 9.9292026 | 68.052348 | 2.17351734 |
| VIT_10s0003g02810 | WRKY transcription factor | 1.36916 | 3.7707 | 1.4684 | 25.9585 | 27.7092 | 4.30239 |
| VIT_01s0026g01730 | WRKY transcription factor | 1.723862 | 14.588 | 2.922619 | 5.46501211 | 27.036 | 2.7636535 |
| VIT_05s0077g00730 | WRKY transcription factor | 13.330992 | 26.9002 | 7.246623 | 75.25308 | 109.90206 | 22.386063 |
| VIT_10s0003g01600 | WRKY transcription factor | 11.8002 | 76.5289 | 17.1084 | 47.3846 | 104.561 | 46.8014 |
| VIT_02s0025g00420 | WRKY transcription factor | 0.195665 | 2.59785 | 0.624419 | 0.464592 | 13.4431 | 1.64438 |
| VIT_11s0052g01620 | Pathogenesis-related protein PR-1 | 2.35673 | 0.90294 | 3.85426 | 0.976269 | 18.1247 | 7.39029 |
| VIT_14s0081g00020 | Pathogenesis-related protein PR-1 | 5.2547252 | 4.563528 | 0.86386283 | 17.212108 | 69.535528 | 4.8717034 |

### *Verification of candidate* WRKY53 *genes*

In higher plants, *WRKY* genes play a variety of roles. Accumulating evidence indicates that WRKY transcription factors are involved in the responses to biotic stresses as well as in plant development (Chen and Chen, 2000; Du and Chen, 2000; Eulgem *et al.*, 2000). WRKY proteins comprise a large family of transcription factors (Ulker and Somssich, 2004) that are potentially involved in the regulation of transcriptional reprogramming responsible for plant immune responses (Eulgem and Somssich, 2007). In our results, eleven *WRKY* genes were identified as candidate resistant genes because their expression was positively correlated with resistance to *C. diplodiella* in *DAC*. One of these WRKY genes, *VdWRKY53* (Genbank accession KY124243) was cloned from *DAC* and further characterized. The WRKY domain of VdWRKY53 belonged to Group III subfamily of WRKY family. In plants, the Group III subfamily of WRKY were considered to be the most evolutionarily advanced and the most adaptable group, and to be co-evolved with disease resistance genes. VdWRKY53 was found to be closely related to VvWRKY30, 46, 41 and AtWRKY41 and 53 in a phylogenetic analysis using all members of WRKY III subfamily from *Vitis* and *A. thaliana* (Fig. 5). *A. thaliana* wth *AtWRKY53* loss of function mutants showed delayed development of disease symptom after infection of *Ralstonia solanacearum*, but increased susceptibility toward *Pseudomonas syringae* (Murray *et al.*, 2007). The expression of AtWRKY41 is specifically suppressed by a compatible strain of *P. syringae* in an effector-dependent manner (Pandey and Somssich, 2009).

**Fig. 5.**
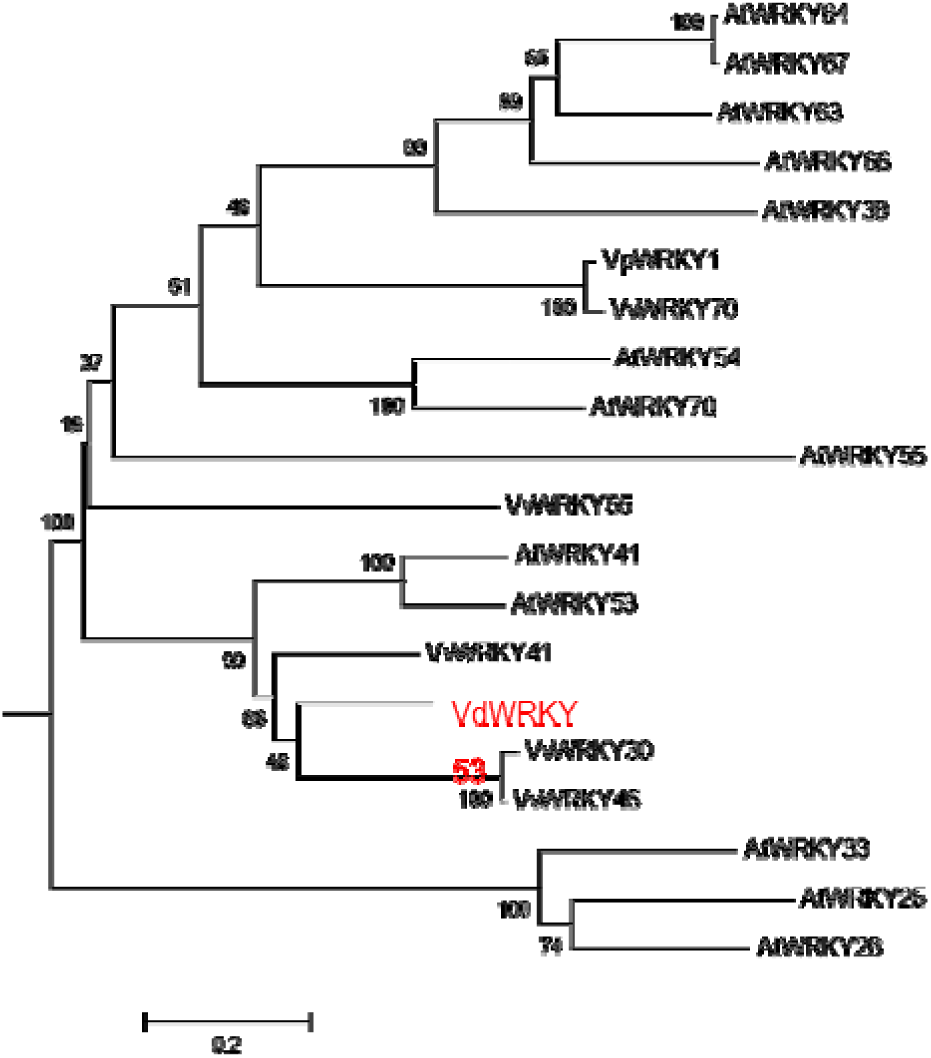
The phylogenetic relationship among the subfamily III WRKY transcription factors of Vitis and *A. thaliana*. The phylogenetic tree were constructed using full-length amino acid sequence (listed in Supplementary 4) and neighbor joining (NJ) method in Clustal X version 1.83 and Mega version 5.0. Bootstrap values (shown on branches) were calculated from 1,000 iterations.

Classifying VdWRKY53 into the same clade as AtWRKY41 and 53 suggests a function of VdWRKY53 in disease resistance. To confirm the function of VvWRKY53, Arabidopsis transgenic plants were produced using the pGW53 construct for over-expression of VvWRKY53. Homozygous transgenic plants were identified from three independent transgenic lines, GW53-1, -2, and -3, by growing them to T3 generation. The homozygous transgenic plants and wild-type Col plants were infected by *C. diplodiella*, *G. cichoracearum* and PDC3000. As determined by q-RT-PCR analyses, *VdWRKY53* was expressed in all three transgenic lines prior and post infection of the three diseases. The expression levels were between 0.5 and 2 folds of the mRNA levels of the Arabidopsis house-keeping gene *AtActin* (Fig 6). Signal for *VvWRKY53* expression in WT Col plants were at background level for most samples, and at a low level for some samples of Col-2. The low level of signal could be the result of non-specific amplification of Arabidopsis genes with homology to the primers. Consistent with the over-expression of *VdWRKY53* in GW53, the disease resistant level was remarkable improved. The GW53 plants were more resistant to *C. diplodiella*, PDC3000 and *G. cichoracearum* than Col plants. After infection by *G. cichoracearum*, GW53 plants grew normally with green and health leaves even powdery mildew was present on their leaves. Whereas, WT Col plants developed a strong disease symptom, with yellow and even dead leaves. The same results happened to *C. diplodiella* and PDC3000 infection. Most of GW53 plants could grow normally with *C. diplodiella* and PDC3000, but Col could not (Fig. 7). For GW53 transgenic plants, 95%, 98% and 100% of leaves were free of disease symptoms after infection with *G. cichoracearum*, *P. syringae* PDC3000, and *C. diplodiella* respectively (Fig. 8). For WT Col plants, only 5%, 0% and 2% leaves were free of disease symptoms after infection with the three diseases respectively.

**Fig. 6.**
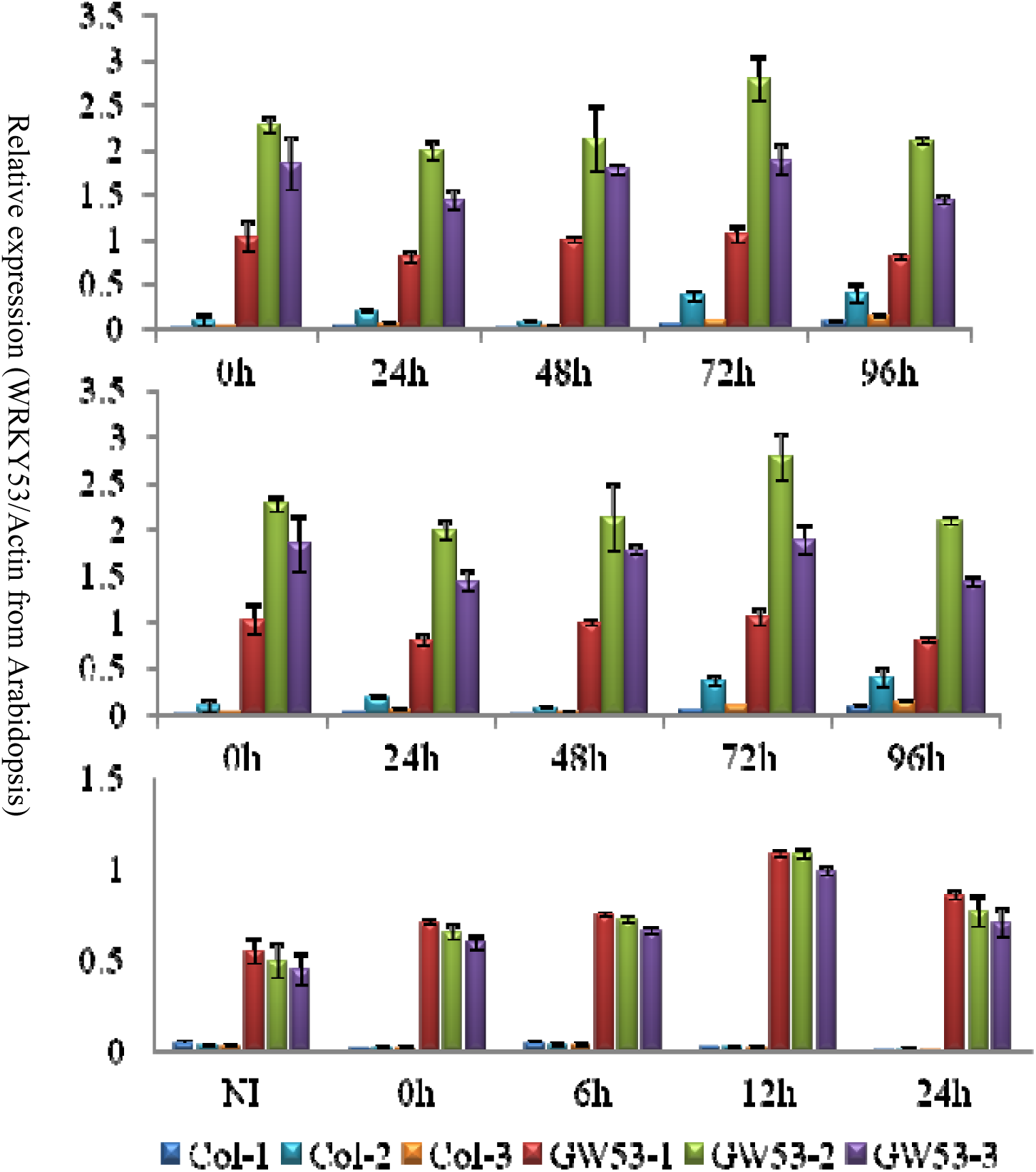
Quantitative PCR analyses of VdWRKY53 gene expression in wildtype and transgenic Arabidopsis plants. The expression levels of *VdWRKY53* in three wildtype (Col-1, 2, 3) and three transgenic lines (GW53-1, -2, and -3) were analyzed at different stages of infection by pathogens *G. cichoracearum*, *C. diplodiella* and *P. syringae* pv *tomato* PDC3000. Error bars represent the standard deviation of three independent PCR reactions. NI, not infected. h, hours after infection.

**Fig. 7.**
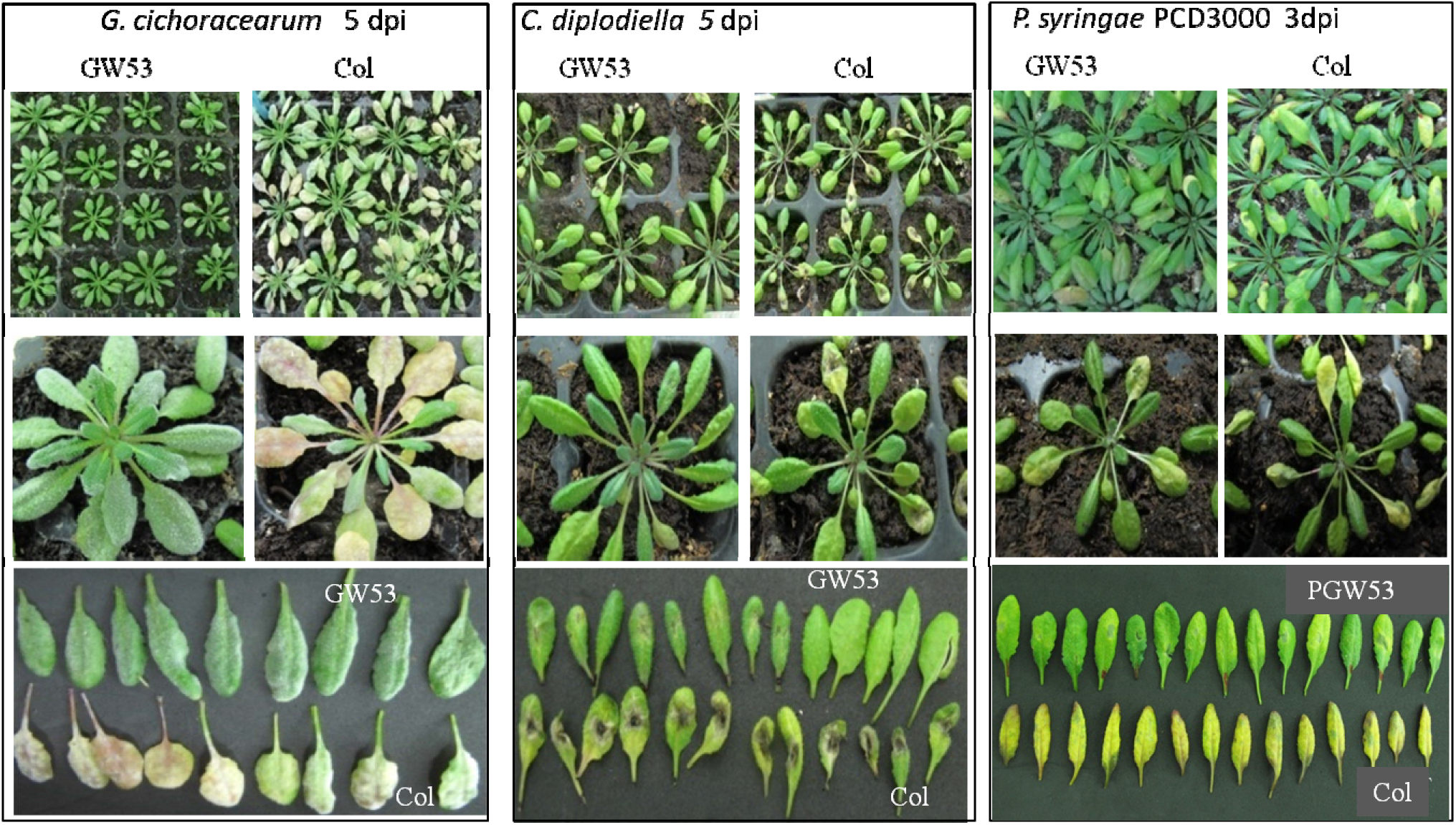
Transgenic *Arabidopsis* plants over-expressing *VdWRKY53* showed enhanced resistance to pathogens. Kanamycin resistant T2 plants from three different T1 lines were infected with pathogens *G. cichoracearum*, *C. diplodiella* and *P. syringae* pv *tomato* PDC3000, separately. The images show the results of one transgenic line along with wildtype controls at 3 or 5 days post infection (dpi). The plants of other transgenic lines displayed similar results.

**Fig. 8.**
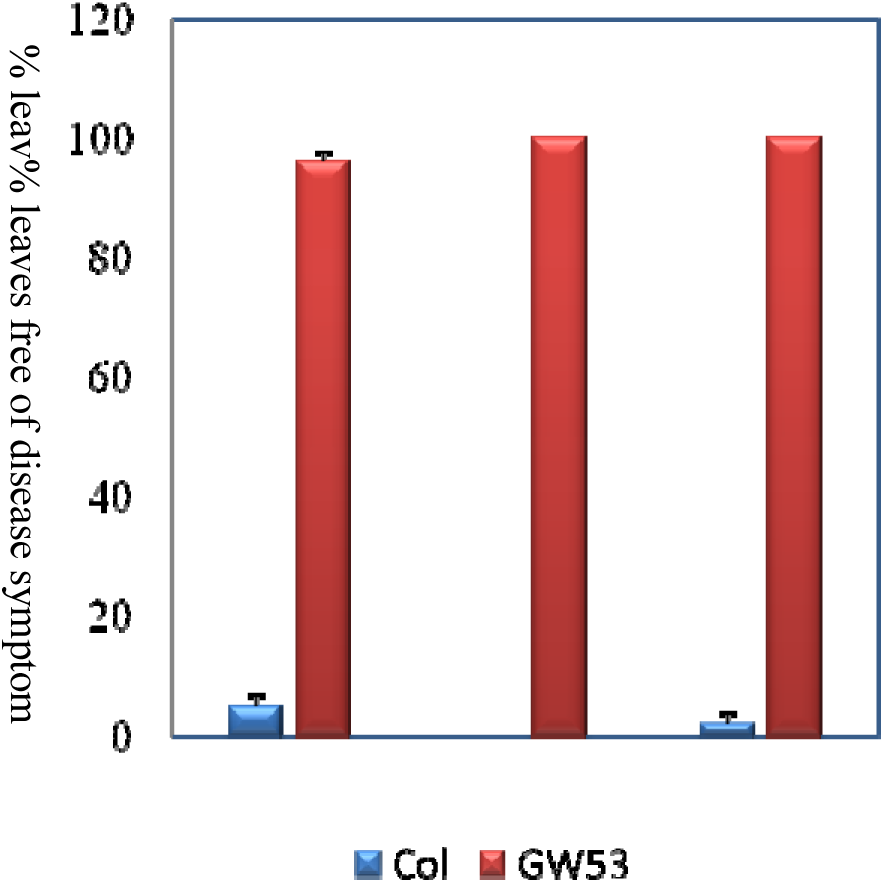
Percentage of Arabidopsis leaves were free of symptoms after pathogen infection. Kanamycin resistant T2 plants from three different T1 lines were infected with pathogens *G. cichoracearum*, *C. diplodiella* and *P. syringae* pv *tomato* PDC3000, separately. Disease symptoms were visually examined on 33 leaves for each line and each infection at 3 dpi (days post infection) for *P. syringae* pv *tomato* PDC3000 and at 5 dpi for *G. cichoracearum* and *C. diplodiella* infection. Error bars represent the standard deviation of three transgenic lines or three populations of wildtype control plants.

## Discussion

### *Rapid response contributes to* DAC *resistance to disease*

The first layer of plant defence against pathogens is the cell wall-associated response: pathogenic microorganisms must actively penetrate the plant apoplast for access. The second layer of plant defence is the HR, in which cell death surrounding an infection restricts the growth of pathogens. Ralph Huckelhoven considered HR-associated cell death to be a complex defence that depends on the timing of HR (Huckelhoven, 2007). In our study, when *C. diplodiella* invaded, *DAC* quickly showed a typical HR quickly. HR and cell death limited pathogen invasion, and then resistance genes in the signalling pathway were switched on (Fig. 1B, C). In comparison with VIM, 20 genes co-expressed in the HR showed changes in *DAC* at the 12 hpi time point (Fig. 2D). This result indicated that a key switch for the *DAC* resistance response occurs near the 12 hpi time point, which was also observed in other resistant grapevine species, such as *Vitis riparia* (resistant) after infection with *Plasmopara viticola* (Polesani *et al.*, 2010). *DAC* could rapidly perceive microbial molecules by surveillance of host cellular intactness, which is a common mechanism in plants. This mechanism has also been observed in *V. riparia* infected with *P. viticola* (Polesani *et al.*, 2010). A special mechanism of pathogen defence was observed in the resistant species *DAC*, which quickly recognized the infection signal and activated the HR reaction.

### *Candidate genes contributed to defence in* DAC

Detailed pathogen resistance mechanisms have been described in plant models. They involve complicated signalling pathways and a cascade of resistance genes triggered by an elicitor. Plants use PAMPs (pathogen-associated molecular patterns) or DAMPs (damage-associated molecular patterns) to recognize general elicitors or a special elicitor, which is similar to the innate immune system in animals (Jones and Dangl, 2006). When a fungus infects plants, there are exchanges of signals between the pathogen and the plants (Grenville-Briggs and van West, 2005).

Cell wall-associated plant defence is the first and most important barrier in basal resistance. Basal resistance seems to be suppressed by virulent pathogens but boosted in induced and race-specific resistance. In *Oryza sativa*, *OsWAK1* transcripts were significantly induced by *Magnaporthe oryzae* and play important roles in rice blast disease resistance (Ali *et al.*, 2009). In this study, two wall-associated receptor kinase (WAK) genes (VIT_18s0041g00020, VIT_18s0001g11620) were listed as candidate resistance genes.

In both plant resistance and animal innate immunity, serine/threonine-rich repeat receptor-like kinases contribute to the detection of conserved nonself molecules (Jones and Takemoto, 2004; Chisholm *et al.*, 2006; Huckelhoven, 2007). Five receptor kinase category candidate genes were listed as candidates, namely, LRR receptor-like serine/threonine protein kinases (VIT_12s0035g00070, VIT_10s0092g00590, VIT_12s0055g00580, Vitis_vinifera_newGene_4892, Vitis_vinifera_newGene_4928). The downstream phosphorylation of a WRKY transcription factor ultimately led to the activation of defence-related genes and the partial restriction of pathogen growth (Asai *et al.*, 2002). WRKY transcription factors are involved in nonspecific fungal and oomycete elicitor signal transduction leading to defence gene expression (Cormack *et al.*, 2002; Huckelhoven, 2007). Eleven WRKY transcription factors contributed to defence in this study, showing co-expression in K03 clusters. WRKY transcription factors ultimately lead to the activation of defence-related genes and the partial restriction of pathogen growth (Ali *et al.*, 2009). Pathogenesis-related proteins are secreted into the defence system during the resistance response; for example, pathogenesis-related protein 1 (PR-1) has antifungal and antibiotic activity (Rauscher *et al.*, 1999; Huckelhoven, 2007; Yu *et al.*, 2013). In our data, two PR-1 protein type genes (VIT_11s0052g01620, VIT_14s0081g00020) in *DAC* were shown to participate in the defence against pathogen invasion. In plants, the resistance response is a very complicated system that includes structure and strengthening the cell walls (Huckelhoven, 2007) and the PR protein HR (Greenberg and Yao, 2004), through which cells initiate death after infection to block further infection by the pathogen and allow the plant to survive.

### *Candidate gene* VdWRKY53 *improved the resistance of* Arabidopsis

WRKY transcription factors comprise a large family of regulatory proteins and have been implicated in the defence against pathogens in plants (Pandey and Somssich, 2009). In grapes, *VvWRKY1* and *VvWRKY2* conferred enhanced resistance against fungal pathogens in transgenic tobacco plants(Guillaumie *et al.*, 2010). *VvWRKY11* provides higher tolerance to water stress induced by mannitol compared with the tolerance in wild-type plants, indicating that *VvWRKY11* is involved in the response to dehydration stress (Liu *et al.*, 2011). The *VpWRKY1*, *VpWRKY2* and *VpWRKY3* genes isolated from *Vitis pseudoreticulata* enhanced resistance to biotic and abiotic stress responses (Zhu *et al.*, 2012). VdWRKY53 belongs to the group III WRKY transcription factors and is very similar to AtWRKY53 (Fig. 4). The *AtWRKY53* gene was rapidly induced under drought conditions (Sun and Yu, 2015) and found to positively regulate the basal resistance to *P. syringae* in combination with the *AtWRKY46* and *AtWRKY70* genes (Hu *et al.*, 2012). SolyWRKY53, an orthologous gene to AtWRKY53, is also resistant to TYLCV infection (Huang *et al.*, 2016). In this study, the *VdWRKY53* gene showed higher expression in *DAC* than in *VIM* even at 0 hpi. The RPKM value reached 109.9 at approximately 12 dpi and contributed to the resistance to *C. diplodiella* (Table 4). We cloned the *VdWRKY53* CDS sequence and transferred it into Arabidopsis to generate five lines of transgenic plants over-expressing *VdWRKY53*. As we expected, *VdWRKY53* conferred strong resistance to *C. diplodiella*, PDC3000 and *G. cichoracearum*. Candidate genes from the K03 cluster were co-expressed and located in the same resistance pathway (Broekaert *et al.*, 2006; Eulgem, 2006; Eulgem and Somssich, 2007; Zhang and Feng, 2014; Ma *et al.*, 2016). In our research, *VdWRKY53* improved the resistance of GW53 to infection with *C. diplodiella*, PDC3000 and *G. cichoracearum* separately.

Innate immune perception triggers both local and systemic responses, allowing a plant to fight off pathogens both in a rapid and localized manner and on an extended scale of time and space. The plant defence response to pathogen invasion involves multiple biological processes: first, recognition of virulence factors; second, transfer to signalling modules (NB-LRR, NLRs and LRR receptor–like kinase); then, regulation of switch genes or proteins (WRKY) by different modules; and finally, activation of the resistance pathway response. In this study, WAK, LRRs, WRKYs, and PRs were in the K03 cluster, and co-expression was induced by the grapevine white rot pathogen in *DAC at* 12hpi time point. The key *WRKY* gene *VdWRKY53* improved Arabidopsis resistance to pathogen and bacteria invasion. Based on these data, we proposed that 20 candidate genes contributed to the resistance of *DAC* to grapevine white rot pathogen.

## Supplementary data

Table S1. Primers of test genes

Table S2. KEGG pathways of DEGs at *V.davidii* and *V.vinifera* different infecting stages

Table S3. Details of KEGG pathways in the research

Table S4. Details of transcripts in k=03

## Acknowledgements

This work was financially supported by the National Natural Science Foundation of China (No. 31201599), the China Agriculture Research System (CARS-30), the Agricultural Science and Technology Innovation Program (CAAS-ASTIP-2017-ZFRI), and the Central Public-Interest Scientific Institution Basal Research Fund (1610192016202, 1610192017202).

